# Behavioral changes preceded by subthalamic nucleus alterations in a progressive macaque model of Parkinson’s disease

**DOI:** 10.1101/2024.01.22.576571

**Authors:** Mathilde Bertrand, Jessy Hugues Dit Ciles, Stephan Chabardes, Nicolas De Leiris, Bruno Baudin, Julien Bastin, Brigitte Piallat

## Abstract

Parkinson’s disease (PD) is diagnosed after motor symptoms appear, although non-motor symptoms emerge years earlier. Following years of pharmacological treatment, high-frequency stimulation (HFS) of the subthalamic nucleus (STN), a key hub in goal-direct behaviors, can be proposed. While HFS-STN reliably improves motor symptoms, it does not specifically address non-motor symptoms. Clarifying how STN dysfunction contributes to non-motor symptoms could thus improve STN stimulation strategies. Here, we longitudinally recorded STN local field potentials in two macaques performing a counter-demanding task during chronic low-dose MPTP treatment. This progressive model, evolving from an asymptomatic stage to motivational, cognitive and ultimately motor deficits, enabled detailed examination of non-motor stages preceding motor impairment. Each stage was associated with distinct STN electrophysiological alterations, including early loss of reward-related theta activity, subsequent disappearance of decision-related theta oscillations, and later reduction of movement-related beta rebound. In the stable parkinsonian stage, stimulation of different STN territories provided complementary behavioral effect: dorsal HFS-STN improved motor performances, whereas ventral low-frequency stimulation alleviated motivational deficits. These findings reveal a temporal link between STN dysfunction and symptom onset, and suggest site and frequency-specific stimulation as a strategy to address both motor and non-motor symptoms in PD.

## INTRODUCTION

Parkinson’s disease (PD) is a progressive neurodegenerative disorder characterized not only by cardinal motor symptoms but also by a range of non-motor symptoms, including sleep/wake disturbances, apathy, and cognitive decline, which precede motor signs by years^1–3^. Understanding the neural substrates of these early manifestations is critical for enabling early diagnosis and developing targeted therapies^4^. The subthalamic nucleus (STN) is a central hub in the basal ganglia-thalamo-cortical network, integrating cortical inputs to shape and adapt goal-directed behaviors^5,6^. In PD, pathological STN beta-band activity (12-35 Hz) is associated with motor symptoms occurrence and abolished by dorsolateral STN high-frequency stimulation (HFS, ∼130 Hz), which improves motor symptoms^7–10^. Beyond motor control, accumulating evidence implicates STN theta-band (4-8 Hz) alterations with non-motor PD symptoms, including impaired conflict resolution, apathy, impulsivity, and depression^11–15^.

We recently demonstrated correlations between physiological STN activity and distinct behavioral domains in non-human primate (NHP), providing a framework for studying how these dynamics progress during PD^16^. Among available preclinical models, the NHP parkinsonian model induced by 1-methyl-4-phenyl-1,2,3,6-tetrahydropyridine (MPTP) remains the gold standard, as it reproduces both motor and non-motor impairments alongside nigro-striatal degeneration^17–19^. In particular, chronic low-dose MPTP treatment induces a progressive PD phenotype^20,21^ enabling longitudinal study of STN activity and its temporal relationship to the onset of non-motor and motor manifestations.

Deep brain stimulation (DBS) provides a causal approach to explore STN functions. While HFS of the dorsolateral STN reliably alleviates motor symptoms, it can exacerbate non-motor impairments^22,23^. Conversely, dorsal low-frequency stimulation (LFS, ∼4-5 Hz) has shown promise for improving non-motor symptoms in PD patients, including deficits in conflict resolution and impulsivity, likely by engaging prefrontal-STN hyperdirect pathway^24–26^. With emerging adaptive DBS technologies enabling site and frequency-specific modulation^27–30^, clarifying the temporal and functional dynamics of STN alterations has become a pressing clinical priority.

Here, we investigated the time-based longitudinal relationship between STN activity and the sequential emergence of motivational, cognitive, and motor deficits in a progressive NHP model of PD. We reveal that STN alterations precede behavioral changes, with theta disruptions related to motivational and cognitive deficits, and abnormal beta activity associated with motor symptoms. In a stable parkinsonian syndrome, we further demonstrate site and frequency-specific STN stimulation effects, with dorsal HFS improving motor impairments and ventral LFS alleviating motivational deficits.

## RESULTS

### Progressive parkinsonian syndrome

Chronic low-dose MPTP administration (0.2-0.5 mg/kg, biweekly) induced a progressive parkinsonian syndrome in both monkeys. M1 received 20 injections (5.45 mg/kg over 43 weeks) and M2 received 18 injections (6.7 mg/kg over 39 weeks), and both reached a similar stable moderate parkinsonian syndrome. Using a validated scale to evaluate motor manifestations^31^, we identified three severity stages (**Fig. 1A**): stage 1 (score 0-3), asymptomatic, with no motor signs; stage 2 (score 4-10), mild parkinsonism, with light and unstable deficits; and stage 3 (score >10), moderate parkinsonism, with significant and stable motor manifestations, including reduced activity, bradykinesia, and flexed posture.

**Figure 1:**
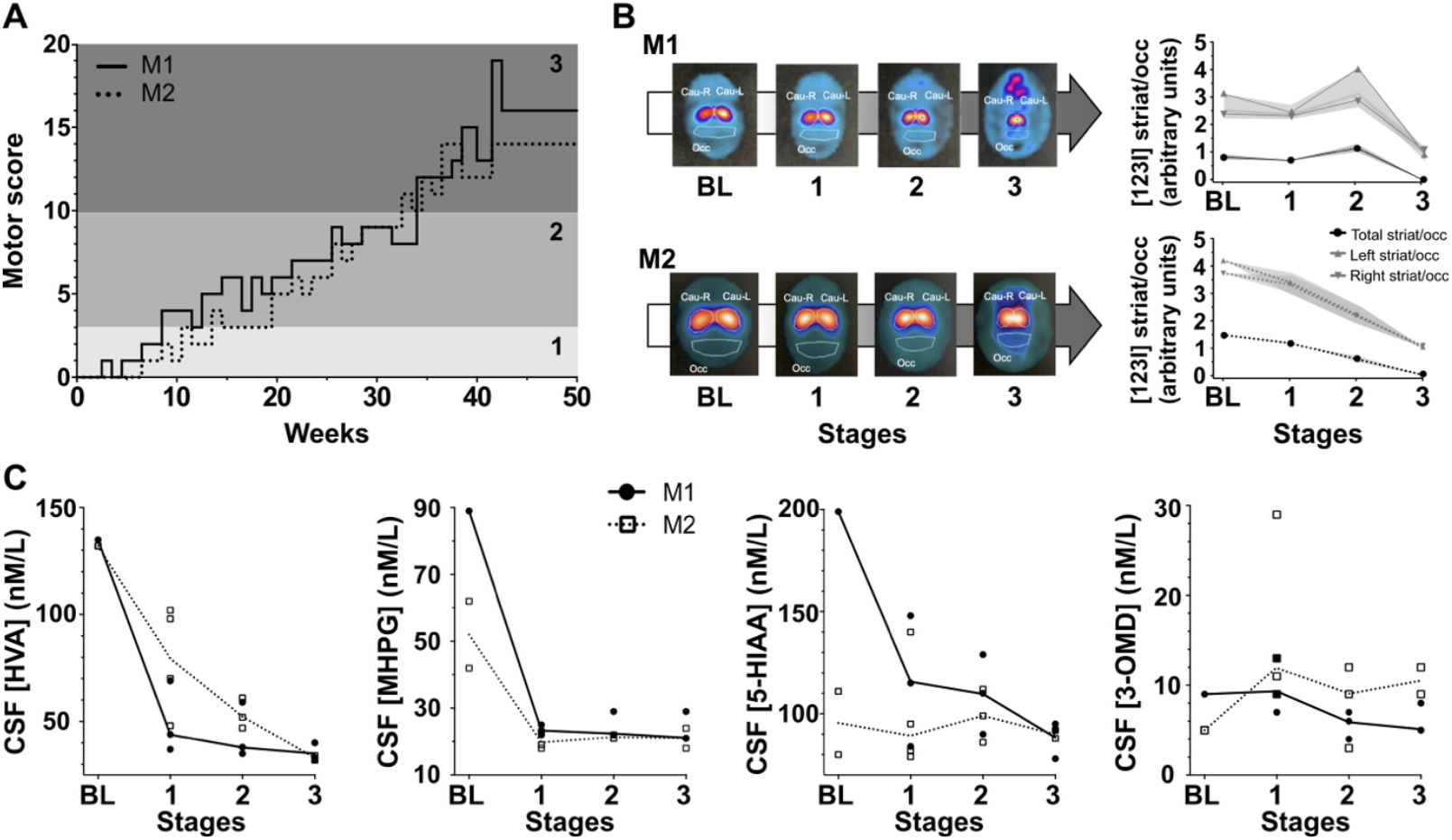
Progressive parkinsonian syndrome for M1 (solid line) and M2 (dotted line). **(A)** Weekly evaluation of the parkinsonian syndrome induced by chronic low-dose MPTP administration. Motor score discriminated 3 stages: 1-asymptomatic (very light gray, score 0-3); 2-mild parkinsonism (light gray, score 4-10); 3-moderate parkinsonism (dark gray, score >10). **(B)** DaT SPECT imaging using Ioflupane (I^123^) tracer for each stage with ratios quantification between the left and right striatum (Cau-L and Cau-R) relative to cerebral background noise (Occ). **(C)** Lumbar cerebrospinal fluid (CSF) concentrations of homovanillic acid (HVA), 5-hydroxyindole acetic acid (5-HIAA), 3-methoxy-4-hydroxyphenylglycol (MHPG), and 3-ortho-methyldopa (3-OMD). BL, baseline before MPTP treatment.

Longitudinal DaT SPECT imaging revealed a progressive decrease in striatal Ioflupane tracer relative to the occipital cortex, beginning after the first MPTP injection (**Fig. 1B**). Both monkeys developed similar trajectories, except for a transient increase during stage 2 for M1, which led to a closer behavioral examination during this stage.

Finally, we analyzed serial cerebrospinal fluid (CSF) to identify metabolite correlates of the dopaminergic deficit (**Fig. 1C**). In both monkeys, homovanillic acid (HVA), principal dopamine metabolite, progressively decrease and 3-methoxy-4-hydroxyphenylglycol (MHPG), major noradrenaline metabolite, was significantly reduced from stage 1. M1 also showed a progressive decrease in 5-hydroxy indoleacetic acid (5-HIAA), main serotonin metabolite, whereas 3-ortho-methyldopa (3-OMD), product of dopamine synthesis, remained unchanged.

### Early non-motor manifestations

Before and during MPTP treatment, both monkeys performed a validated counter-demanding task discriminating motivational, cognitive and motor behaviors^16^. On each trial, they could either perform a switching task (Work), where correct responses filled a gauge, or checked the gauge itself (Check) to monitor their progress toward a bonus reward delivered once full (**Fig. 2A**, see methods). Motivation was assessed by the probability to check when the gauge was full (Reward Check); cognitive performance by the error rate on Switch trials (Error Switch) and Switch-Cost (response time difference between correct Switch and Non-Switch trials); and motor performance by response time on correct Non-Switch trials (mRT). Each monkey completed at least five sessions per stage (≥4 bonus rewards/session; **Supplemental Table 1**): M1 performed 10, 17, 13, 6 and 6 sessions at baseline, stage 1, stage 2, stage 2’, and stage 3, respectively; and M2 performed 10, 16, 9, 7, and 5 sessions.

**Figure 2:**
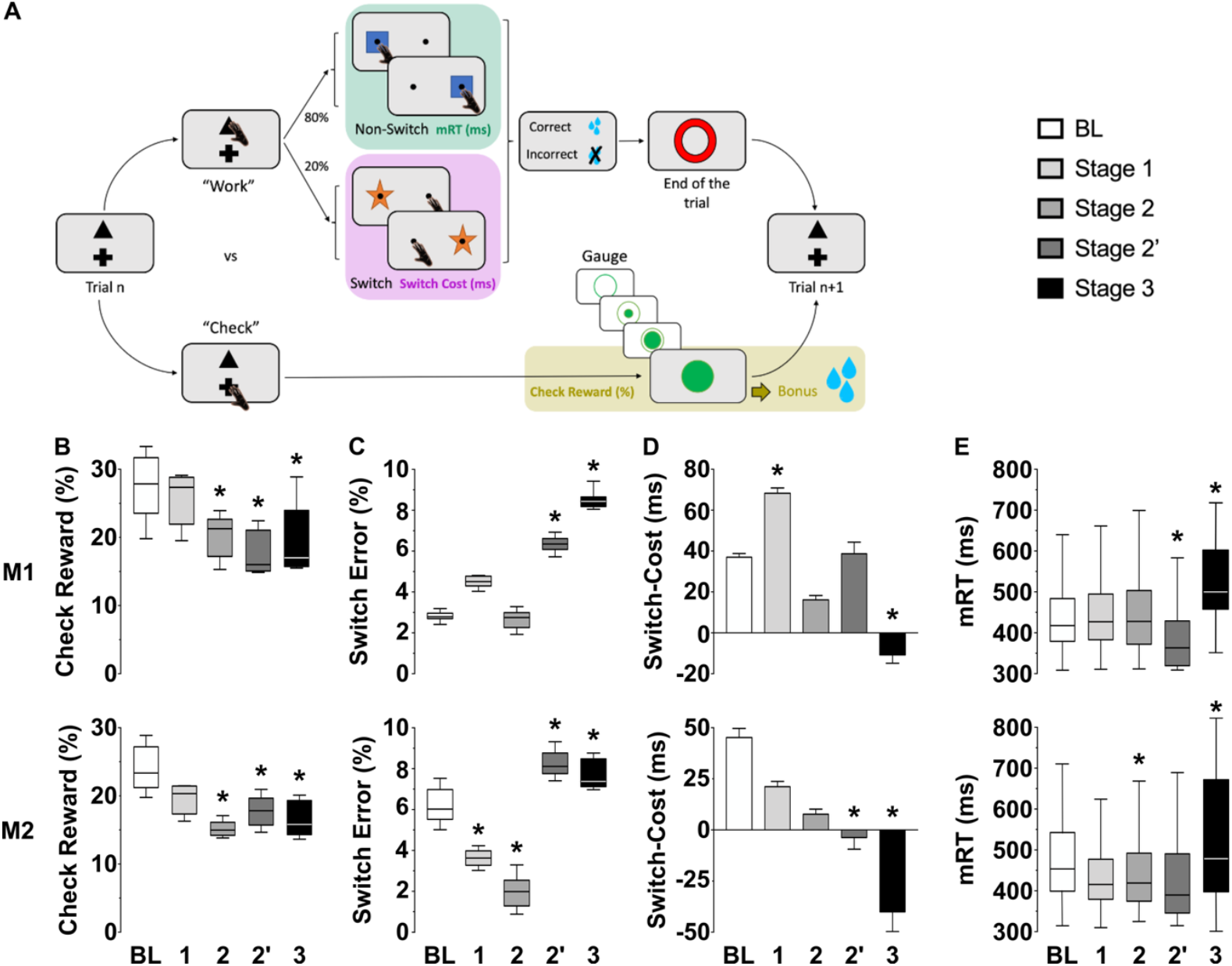
Behavioral effect of MPTP treatment across stages. **(A)** Rewarded task assessing limbic (gold), cognitive (pink), and motor (green) behaviors. On each trial, monkeys decided to “Work” on the main switching task, or “Check” the size of a gauge indicating the proximity of a bonus reward (see Methods). **(B)** Probabilities to check when the reward is available (Check Reward, %). **(C)** Errors on Switch trials (Switch Error, %). **(D)** Switch-Cost in ms, response time difference between Switch and Non-Switch trials. **(E)** Motor response time (mRT, ms) on Non-Switch trials. Stages: baseline, before MPTP treatment (BL, white); stage 1, asymptomatic (1, very light gray); early stage 2, mild parkinsonism (2, light gray); late stage 2’, mild parkinsonism (2’, light gray), stage 3, moderate parkinsonism (3, dark gray). The horizontal line represents the median, except for Switch-Cost (mean ± SEM). *p<0.05, Kruskal-Wallis followed by Dunn’s multiple comparisons test.

During stage 1, just after starting MPTP treatment, no significant behavioral effects were found for both monkeys, consistent with an asymptomatic stage. In stage 2, non-motor signs appeared in a specific chronology, distinguishing two substages: early stage 2, affecting motivation with decreased Check Reward medians (for **M1-** BL: 28% (IQR 23-32) vs. stage 2: 21% (17-23), p=0.039; for **M2-** BL: 23% (21-27) vs. stage 2: 15% (14-16), p=0.010) (**Fig. 2B**); and late stage 2’, affecting cognitive behavior with increased Switch Error (for **M1-** BL: 2.8% (2.7-3) vs. stage 2’: 6.4% (6-6.6), p<0.001; for **M2-** BL: 6.0% (5.5-7) vs. stage 2’: 8.1% (7.8-8.8), p<0.001) (**Fig. 2C**). M2 also exhibited a mean negative Switch-Cost (BL: 45.1 ± 4.4ms vs. stage 2’: -3.9 ± 5.7ms, p<0.001), suggesting advanced cognitive impairment (**Fig. 2D**).

Finally, during stage 3, both monkeys displayed slower motor response time with increased mRT (for **M1-** BL: 411ms (IQR 372-478) vs. stage 3: 501ms (460-603), p<0.001; for **M2-** BL: 454ms (390-536) vs. stage 3: 487ms (399-676), p<0.001) (**Fig. 2E**).

### Evolution of STN activity

STN local field potentials were recorded while monkeys performed the behavioral task at baseline and throughout MPTP treatment. Time-frequency representations of spectral power were averaged across trials within each condition and aligned to task events. Band-specific powers within defined time windows were then quantified and compared across stages.

During reward delivery, baseline activity showed a significant decrease in theta oscillations (4-8Hz) for both monkeys (p<0.05, two-sample t-test) (**Fig. 3A**). This decreased theta disappeared, starting from stage 1, with theta power significantly higher at all parkinsonian stages compared to baseline (for **M1-** BL: -3.4 ± 0.2LdB vs. stage 1: 1.3 ± 0.1 LdB, p<0.001; for **M2-** BL: -2.2 ± 0.1LdB vs. stage 1: 0.5 ± 0.2LdB, p<0.001).

**Figure 3:**
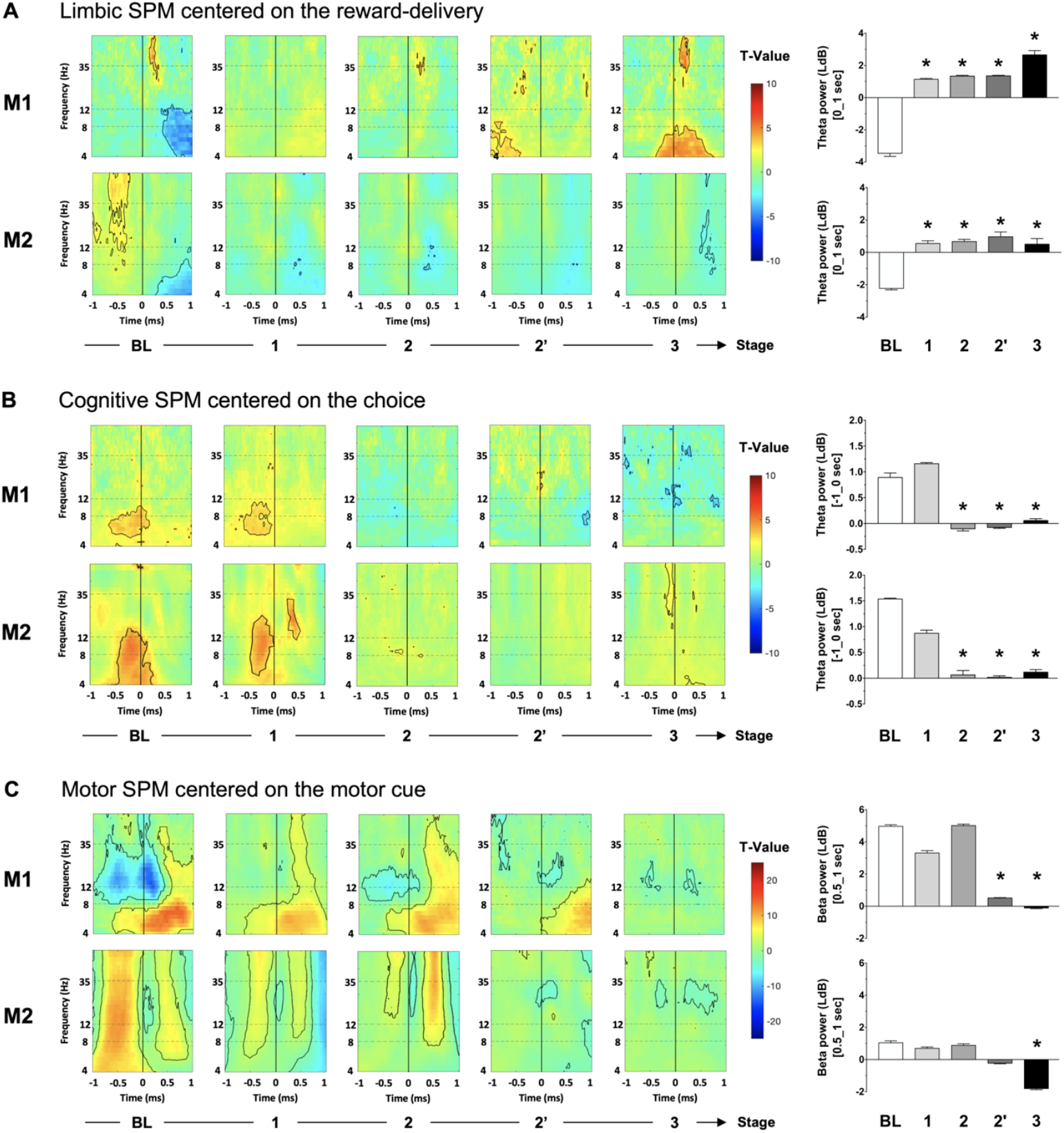
STN neuronal effect of MPTP treatment across stages. **(A)** Limbic statistical parametric maps (SPM) centered on the full gauge with reward delivery (vertical line at 0) relative to the gauge view without the reward with theta quantification between 0 and 1 second. **(B)** Cognitive SPM centered on cognitive choice (vertical line at 0) to work relative to check with theta quantification between -1 and 0 second. **(C)** Motor SPM centered on the onset of Non-Switch stimulus (vertical line at 0) relative to a resting state, with beta quantification between 0.5 and 1 second. Stages: baseline, before MPTP treatment (BL); 1, asymptomatic; 2, early mild parkinsonism; 2’, late mild parkinsonism, 3, moderate parkinsonism. Significant values are encircled with solid lines p<0.05. Power quantifications are expressed in LdB with mean ± SEM, and *p<0.05.

When monkeys chose to work rather than check, baseline activity showed a significant increase in theta oscillations for both monkeys (p<0.05, two-sample t-test) (**Fig. 3B**). This effect was preserved in stage 1 but disappeared from early stage 2, with theta power significantly lower than baseline (for **M1-** BL: 0.9 ± 0.1LdB vs. stage 2: -0.1 ± 0.1LdB, p<0.001; for **M2-** BL: 1.5 ± 0.1LdB vs. stage 2: 0.1 ± 0.1LdB, p=0.004).

At the onset of the motor cue, baseline activity showed a significant decrease in beta oscillations followed by a rebound beta increase for both monkeys (p<0.05, one-sample t-test) (**Fig. 3C**). This rebound persisted during stage 1 and early stage 2 but disappeared from stage 2’. Quantification of beta rebound confirmed significant reductions at stage 2’ for M1 and at stage 3 for M2 (for **M1-** BL: 5.0 ± 0.1LdB vs. stage 2’: 0.5 ± 0.3LdB, p<0.001; for **M2-** BL: 1.0 ± 0.1% vs. stage 3: -1.8 ± 0.1LdB, p=0.021).

### STN modulation on parkinsonian syndrome

In stage 3, when both monkeys exhibited a stable moderate parkinsonian syndrome, LFS (4 Hz) and HFS (130 Hz) were applied to the ventral and dorsal STN. Each animal completed at least five sessions per condition (≥4 bonus rewards/session; **Supplemental Table 2**). Despite inter-individual variability, consistent with PD patients reports ^32^, both monkeys showed converging effects.

Ventral LFS-STN restored the decreased motivation (Check Reward) to baseline value (for **M1-** MPTP ventral LFS: 27% (IQR 24-27) vs. MPTP: 17% (16-24) p=0.049 and BL: 28% (24-32), p=0.999; for **M2-** MPTP ventral LFS: 22% (21-25) vs. MPTP: 16% (14-19) p=0.044 and BL: 23% (21-27), p=0.998) (**Fig. 4A**).

**Figure 4:**
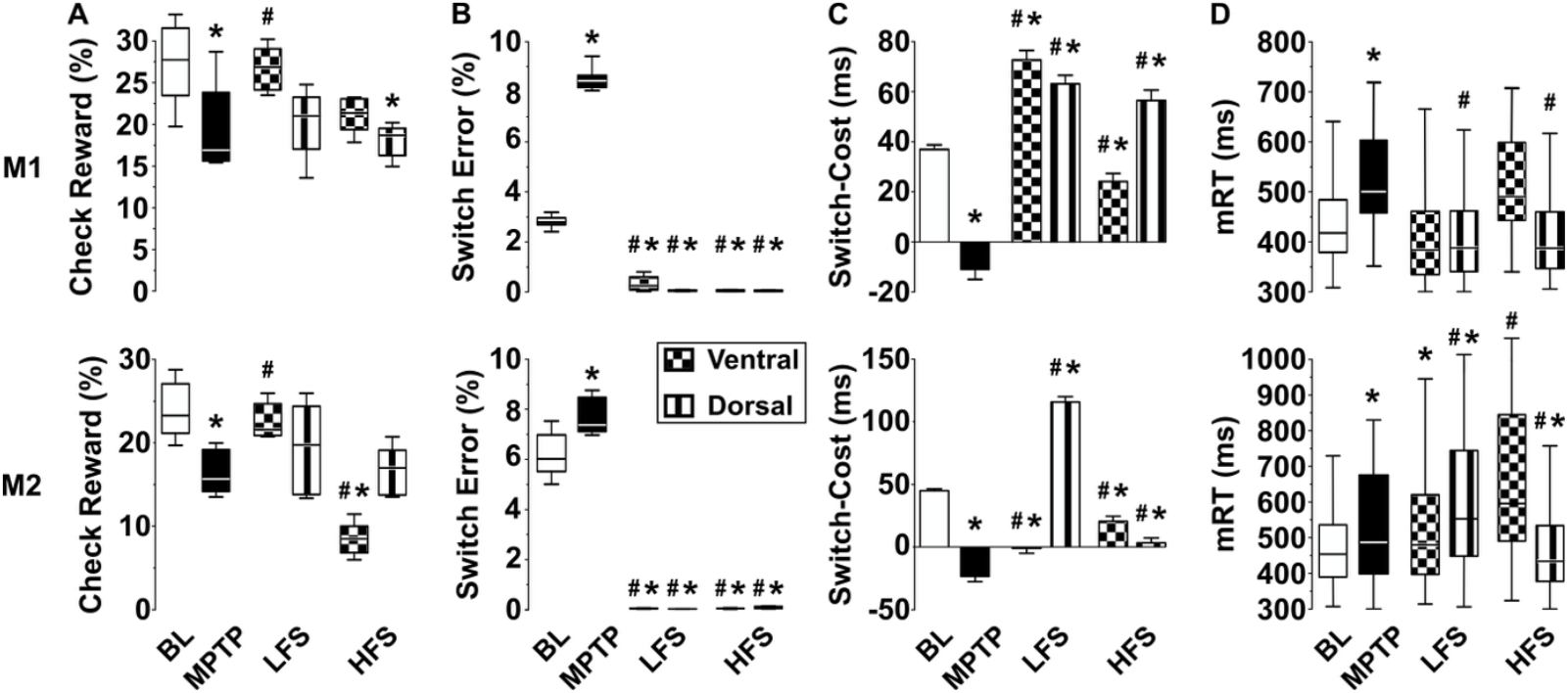
Behavioral effect of LFS (4Hz) and HFS (130Hz) on ventral (grid pattern) and dorsal (vertical lines) STN, on MPTP-treated monkeys. **(A)** Probabilities to check when the reward is available (Check Reward, %). **(B)** Errors on Switch trials (Switch Error, %). **(C)** Switch-Cost in ms, response time difference between Switch and Non-Switch trials. **(D)** Motor response time (mRT, ms). BL, baseline (white); MPTP, stage 3 (black); LFS, 4 Hz applied in stage 3; HFS, 130 Hz applied in stage 3. The horizontal line represents the median, except for Switch-Cost (mean ± SEM). *p<0.05 different from baseline and #p<0.05 different from MPTP off-stimulation, Kruskal-Wallis followed by Dunn’s multiple comparisons test.

Errors on Switch trials were almost eliminated with both LFS and HFS, regardless of dorsal or ventral STN stimulation (for **M1-** MPTP: 8.4% (IQR 8.2-8.7) vs. MPTP ventral LFS: 0.29% (0.09-0.65); MPTP dorsal LFS: 0.06% (0.04-0.08); MPTP ventral HFS: 0.07% (0.03-0.09); MPTP dorsal HFS: 0.06% (0.03-0.06), p<0.001; for **M2-** MPTP: 7.4% (7.1-8.5) vs. MPTP ventral LFS: 0.04% (0.04-0.05); MPTP dorsal LFS: 0.07% (0.04-0.08); MPTP ventral HFS: 0.14% (0.11-0.15); MPTP dorsal HFS: 0.06 (0.06-0.08), p<0.001) (**Fig. 4B**).

In contrast, dorsal LFS-STN increased Switch-Cost compared to both baseline and MPTP (for **M1-** MPTP dorsal LFS: 63.5 ± 3.4ms vs. MPTP: -10.6 ± 4.1ms and BL: 37.0 ± 1.8ms, p<0.001; for **M2-** MPTP dorsal LFS: 116.0 ± 4.3ms vs. MPTP: -40.3 ± 10.6ms and BL: 45.1 ± 1.5, p<0.001) (**Fig. 4C**).

Finally, only dorsal HFS-STN restored the mRT to baseline value (for **M1-** MPTP dorsal HFS: 388ms (IQR 347-460) vs. MPTP: 501ms (460-603), p<0.001 and BL: 411 (372-478), p>0.999; for **M2-** MPTP dorsal HFS: 435ms (378-534) vs. MPTP: 487ms (399-676), p<0.001 and BL: 454ms (390-536), p>0.999) (**Fig. 4D**).

## DISCUSSION

Using chronic low-dose MPTP to develop a progressive PD model, we demonstrated that non-motor impairments emerge prior to motor signs and that these behavioral changes correlate with early alterations in STN activity. In stable parkinsonism, we further show a novel positive effect of ventral LFS-STN in alleviating motivational deficits, while confirming the efficacy of dorsal HFS-STN in managing motor impairments, underscoring the site and frequency-specific contributions of STN circuits to both motor and non-motor outcomes.

### Progressive parkinsonian syndrome

Among NHP models, MPTP uniquely reproduces a comprehensive nigrostriatal dopamine deficit^17^, with human-like clinical features including motor symptoms, sleep/wake disturbances, and cognitive impairments^20,31,33^. In our progressive model, both monkeys developed a months-long worsening parkinsonian syndrome, mirroring the slow progression of human PD and allowing severity-stage discrimination consistent with previous studies. Interestingly, we observed a transient increase in striatal presynaptic dopamine reuptake for M1 during mild parkinsonism. Such fluctuations are consistent with preclinical and clinical observations^36–38^, where increased dopamine synthesis and turnover, as well as downregulation of DaT density, likely contribute to compensatory mechanisms during the prodromal stage.

We also observed CSF decreases in HVA, MHGP, and 5-HIAA, indicating impaired dopamine, noradrenergic and serotonin neurotransmission, patterns reported across PD models and patients^42–44^, and correlating with movement disorders, depression, impulsivity, and cognitive impairments^42,45–47^.

### Behavioral impairments follow STN activity changes

In line with recent studies linking neural activity to PD symptoms severity^48–50^, we demonstrated that alterations in STN activity precede behavioral changes in a defined temporal sequence (motivational, followed by cognitive, and ultimately motor deficits), thereby defining parkinsonian stages **(Fig. 5)**.

**Figure 5.**
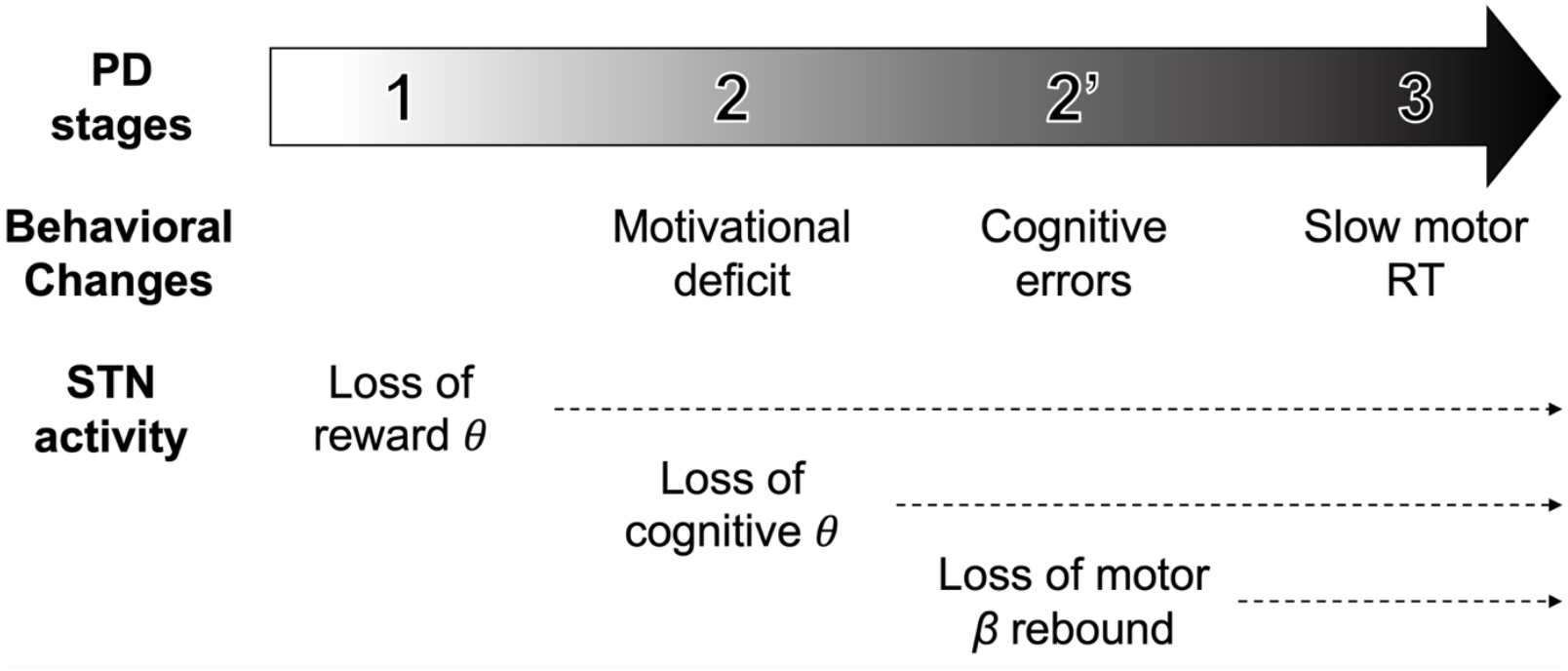
Study-based correlation between subthalamic nucleus (STN) alterations and the subsequent emergence of behavioral changes in a progressive model of Parkinson’s disease (PD). Chronic low-dose MPTP induced distinct stages: 1, asymptomatic; 2, mild parkinsonism with motivational deficits; 2’, novel intermediate stage with cognitive impairments; and 3: moderate parkinsonism with significant motor signs. RT, response time; *θ*, theta activity; *β*, beta activity.

The first stage, after first MPTP injections, is characterized by the absence of significant behavioral changes and therefore considered asymptomatic. In parallel, we observed a baseline-related decrease in theta oscillations during reward delivery, which disappeared from this stage forward. STN theta activity has been associated with limbic processes: in parkinsonian rats as a predictor of addiction vulnerability^51^, and in PD patients with emotional voice processing^52^, reward and loss anticipation^12^, pathological gambling^53^, and depressive symptoms^13^. Notably, an EEG study reported that apathetic PD patients displayed higher baseline alpha-theta power than non-apathetic patients, suggesting that elevated alpha-theta activity may interfere with the neural dynamics required for incentivized movements^54^.

Early stage 2 was defined by reduced motivation, resembling apathy reported in PD models^55,56^ and PD patients^57^. In parallel, decision-related STN theta activity disappeared from this stage onward, consistent with the role of prefrontal-STN circuit in deliberative action selection during a conflict state^11,58^. Additionally, studies have shown that increased STN theta-alpha power is associated with decision-making and conflict resolution^59,60^, whereas a decrease is associated with impulse control deficits and obsessive-compulsive disorder^61,62^.

Late stage 2’ was characterized by cognitive deficits resembling PD mild cognitive impairment^63–65^. Interestingly, both error rate and switch-cost were impacted for M2, whereas only error rate increased for M1, consistent with the transient striatal DaT increase and potential compensatory mechanisms previously discussed. In parallel, movement-related beta dynamics changed: the characteristic movement pattern of beta desynchronization followed by a rebound^66^ was lost from this stage onward. Increased STN beta activity is a validated biomarker in PD and scales with motor symptoms severity^10,67–70^. We therefore hypothesize that motor beta activity variations are flattened by the overall beta increase as parkinsonism progresses.

In stage 3, in addition to motivational and cognitive deficits, clear motor manifestations appeared. These observations are consistent with the course of PD, in which non-motor symptoms precede motor symptoms by several years^2,47^.

### STN modulation

STN-DBS remains the main symptomatic surgical treatment in PD, typically with HFS (130 Hz) targeting dorsolateral STN^7^. Alternative paradigms have been investigated, with dorsal LFS (4-5 Hz) showing promising cognitive improvement^24,25^. Based on these findings, we tested both HFS and LFS in dorsal and ventral STN. Despite inter-individual variability, consistent with clinical reports^27,32^, we observed converging site and frequency-specific effects.

As expected, HFS in dorsal STN improved motor impairments. However, when applied in ventral STN, it aggravated motivational deficits for M2, mirroring the exacerbation of apathy and mood disturbances reported in some PD patients after DBS surgery^22^.

Conversely, LFS in dorsal STN increased the Switch-Cost, consistent with previous 4 Hz STN-DBS findings showing improved inhibition of impulsive/premature responses by engaging prefrontal-STN circuits^24^.

Most strikingly, we found for the first time that ventral LFS restored parkinsonian motivational deficits to baseline levels. This aligns with clinical observations where stimulation closer to the ventral STN is associated with greater improvements in anxiety and depression symptoms^71^; although ventral 4 Hz-LFS was not explored^72^.

### Limitations

While our parkinsonian model offers strong translational value by reproducing a progressive syndrome with both motor and non-motor clinical features, the underlying degenerative mechanisms differ between MPTP toxicity and human PD. Numerous studies have discussed strengths and limitations of this model, most notably the lack of Lewy bodies, a hallmark of alpha-synucleinopathies^17,73–75^.

Another limitation is the small sample size, inherent to NHP research where ethical and logistical constraints preclude large cohorts^76,77^. Nevertheless, intra-individual longitudinal comparisons across parkinsonian stages are a strength of preclinical studies, inaccessible in human research. Finally, variability in STN recording and stimulation sites between the two monkeys may have contributed to inter-individual discrepancies, similar to clinical DBS where subtle differences in lead location strongly influence behavioral outcomes^78,79^.

### Conclusion

Our findings demonstrate that progressive dopamine depletion reshapes STN functional activity in a way that predicts the emergence of non-motor and motor manifestations. Motivational and cognitive impairments should therefore be expected early in PD and actively assessed with appropriate clinical evaluations. Furthermore, the site and frequency-specific effects of ventral LFS and dorsal HFS highlight the therapeutic potential of adaptive DBS strategies, offering a promising approach to manage both motor and non-motor symptoms in PD.

## MATERIAL AND METHODS

### Animals

This study involved two 8 years old monkeys (*Macaca fascicularis*, CRP, Port Louis, Mauritius), one male (M1, 9 kg) and one female (M2, 4 kg). They were pair-housed in a temperature (22 ± 1°C) and humidity (50 ± 5%) controlled facility with a 12h light-dark cycle. They had free access to primate chow and water, and supplemental fruits were given once a day. All procedures followed the European Communities Council Directive of 2010 (2010/63/UE) for care of laboratory animals with the recommendations of the French National Committee (2013/113) and were approved by the local Ethical Committee (#04; authorization n°2019013116115695).

### Experimental design

#### Behavioral task

Monkeys performed a counter-demanding task assessing limbic, cognitive and motor behaviors, as previously described ^16^. Animals were habituated to a primate chair (Crist Instrument Co., MD, USA) and interacted with a touch screen (Elo Touch Solutions, Inc.). In each trial, they chose either to Work on rewarded switching task (triangle, upper center) or to Check their progress with a gauge indicting the proximity of a bonus reward (cross, bottom center). When selecting Work, a blue square (Non-Switch) or an orange star (Switch) appeared randomly left or right and monkeys had to touch the blue square directly or the opposite side of the orange star. Non-Switch trials occurred with higher probability (80%) than Switch trials (20%). Correct responses within 2000 ms were rewarded (sweet liquid, 1 ml) by a computer-controlled system (Crist Instrument Co.), and each trial ended with a 1000 ms red circle before the next Work/Check choice. In Check trials, a gauge (large green circle) was displayed for 5000 ms, filling proportionally with correct Work responses (randomly 8, 16, 24, or 32 required). When Checking a full gauge, a bonus reward (sweet liquid, 5 ml) was delivered, after which the gauge reset. Incorrect responses did not affect the gauge.

#### Behavioral analyses

Only sessions with more than four earned bonus rewards were included. Response times (RT, time between the appearance and touch of the stimulus) were measured from correct trials: motor RT (mRT, motor behavior) for Non-Switch trials and cognitive RT (cRT) for Switch trials. Differences between cRT and mRT represented the Switch-Cost (cognitive behavior). Error rate on Switch trials (cognitive behavior) and probabilities to Check when the gauge was full (Reward Check, limbic behavior) were calculated.

### Parkinsonian syndrome

#### MPTP treatment

Monkeys were chronically intoxicated with biweekly intramuscular 1-methyl-4-phenyl-1,2,3,6-tetrahydropyridine (MPTP) injections (0.2-0.5 mg/kg, in NaCl 0.9%) under light anesthesia (Ketamine 2-4 mg/kg), as previously described ^31^. After each injection, motor score was assessed, and doses were adjusted. M1 received 20 injections over 43 weeks (5.45 mg/kg total) and M2 received 18 injections over 39 weeks (6.7 mg/kg total). Both exhibited a similar stable parkinsonian syndrome.

#### Motor score

Behavioral evaluation was performed, throughout MPTP treatment in the home cage, using a validated macaque parkinsonism scale (Imbert et al., 2000; Davin et al., 2022). Eight criteria were scored from 0 (no impairment) to 3 (maximal impairment), including each arm movement frequency, general activity, posture, bradykinesia, tremor, feeding, freezing and vocalizations (**Supplemental Fig. 1**). According to the literature, three severity stages were identified: stage 1, asymptomatic (0-3); stage 2, mild parkinsonism (4-10); and stage 3, moderate parkinsonism without care needed (>10) (Devergnas et al., 2014; Davin et al., 2022).

#### DaT SPECT imaging

Striatal dopaminergic system was assessed at baseline and across parkinsonian stages using Ioflupane tracer (I123, 90 MBq), which binds to presynaptic dopamine transporters (DaT). For all exams, monkeys were anesthetized for intravenous tracer injection, and images were acquired 3 hours later. A zoom lens was applied for M2 due to sexual dimorphism. Semi-quantitative analysis of striatal ROI was calculated from the three slices with maximal radioactivity and normalized to an occipital cortex ROI.

#### CSF analysis

At baseline and throughout MPTP treatment, CSF was collected under anesthesia by lumbar puncture (L4/L5) using a 22 g Quincke spinal needle (BD™). Sample were stored on ice in sterile 1.5 mL polypropylene tubes, and frozen at -80°C. Monoaminergic metabolites (homovanillic acid, HVA; 5-hydroxyindole acetic acid, 5-HIAA; 3-methoxy-4-hydroxyphenylglycol, MHPG; and 3-orthomethyl-DOPA, 3-OMD), were quantified by ion pair high performance liquid chromatography with electrochemical detection, as previously described ^81^.

### Surgical procedure

Electrode implantation was performed under aseptic conditions and general anesthesia, as previously described ^16^. Monkeys were initially anesthetized with ketamine (7 mg/kg) and xylazine (0.6 mg/kg), and maintained anesthetized with isoflurane. Lidocaine 1% was used for local anesthesia. Saline solution (NaCl 0.9%, Sigma-Aldrich) was infused intravenously during surgery for drug access and hydration. Pre-operative 7T MRI (IRMaGE, Grenoble) allowed the STN identification and per-operative X-rays with ventriculography localized the anterior and posterior commissures (ACPC), with 2ml injection of contrast agent (Iopamiron 200, iodine 200 mg/ml, Bracc) into the left lateral ventricle. STN borders were refined using micro-recordings (250 µm diameter, 0.8-1.2 MOhm, FHC), based on firing rate and pattern. The right STN was implanted with a quadripolar DBS electrode (0.8 mm diameter, four 0.5 mm contacts in two rows, 0.5 mm apart, and and two columns; Heraus©, USA), allowing recordings and two-level stimulations. Electrode was placed at the following stereotactic coordinates: anterior 6/12^th^ of ACPC, 4–5 mm from the midline, with deepest contacts at the lower STN border. The implantation was confirmed by radiography and ventriculography (see ^16^, **Fig. 1D-E**), and the reference fixed at the left occipital level. A stainless-steel head holder (Crist Instruments, MD) was placed at the back of the skull to maintain the head. Post-operative analgesic/anti-inflammatory therapy (Ketoprofen 2mg/kg) were administrated for one week.

### STN recording

Baseline recordings began 2 weeks after surgery and continue throughout MPTP treatment, using a common occipital reference and a multichannel system (AlphaOmega Engineering, Israel). Signals were sampled at 1375 Hz, aligned to task events (± 1000 ms) and analyzed using MATLAB and ImaGIN toolbox (MathWorks). Time-frequency representations of spectral power (2-200 Hz) were computed using a multitaper sliding window with orthogonal discrete prolate Slepian spheroidal (DPSS) tapers, adapted to the frequency and sliding-window length. Single-trial power spectra were averaged by condition and normalized to a pre-stimulus baseline (-1000 to -250 ms) using the formula LdB = 10 log_10_ (P/B), where P is peristimulus power and B is baseline power. Data are expressed in LdB (mean ± SEM) for specific frequency bands defined as theta (4-8 Hz), alpha (8-12 Hz), beta (12-35 Hz), and gamma (35-200 Hz).

### STN stimulation

After MPTP treatment, continuous bipolar stimulation was applied in ventral or dorsal STN, at low frequency stimulation (LFS, 4 Hz), shown to improve cognitive deficits in PD patients ^24,25^, and high-frequency stimulation (HFS, 130 Hz), clinical standard for PD motor symptoms ^7^. For each frequency, stimulation ranges were conducted until side effects occurred (e.g., ipsilateral monocular deviation, head rotation, lip contraction), and intensities set at 80% of this threshold (0.08-0.2 mA; pulse width 60 µs). The number of trials per condition is detailed in **Supplemental Table 2**.

### Statistical analyses

#### Behavioral data

Statistical analyses were performed with GraphPad Prism 8 (GraphPad Software Inc., USA). After normality testing with Shapiro-Wilk test, Kruskal-Wallis tests, followed by Dunn’s multiple comparisons tests, were applied for all behavioral data (mRT, cRT; Switch-Cost, Check Reward), enabling comparison across stages (**Supplemental Table 1**) and stimulation conditions (**Supplemental Table 2**). Data are presented as median (IQR), except for Switch-Cost (mean ± SEM), and a difference was considered statistically significant for a value of p <0.05.

#### Statistical parametric maps

Post-hoc t-tests for time-frequency representations of spectral power were Bonferroni corrected, with “one-sample” t-test to compare signal of one event, and “two-sample” t-test to discriminate significant differences between two events (T-Values). These analyses were plotted as statistical parametric maps (SPM) with significant value encircled (p <0.05). For band-specific analyses, mean relative power were compared across stages using ANOVA followed by Tukey’s test. Data are presented as mean ± SEM and a difference was considered statistically significant for a value of p <0.05.

## Acknowledgements

This work was supported by the French Agence Nationale de la Recherche (ANR), under grant ANR-18-BS01-0005.

## Data Availability

Data and code are available from the corresponding author upon request.

