## Supplemental Fig.1 for "Behavioral changes preceded by subthalamic nucleus alterations in a progressive macaque model of Parkinson’s disease"

Date: ……………………..

MPTP n° …….
Dose: …………….
Dose cumulative:…..

| Frequency of arm movements (for each arm) | General activity |
| --- | --- |
| 0= no difference with control | 0= no difference with control |
| 1= frequent | 1= frequent |
| 2= sometimes | 2= sometimes |
| 3= none | 3= none |
| **Posture** | **Bradykinesia** |
| 0= normal rising position | 0= no difference with control |
| 1= flexed posture (0°<F<45°) | 1= mild slowing of overall movements |
| 2= severely flexed posture (90°<F) | 2= severe slowing of movements |
| 3= dystonic posture | 3= akinesia (no movements) |
| **Tremor** | **Eating** |
| 0= none | 0= normal eating |
| 1= some episodes | 1= slight decrease |
| 2= frequent episodes | 2= severe decrease |
| 3= persistent | 3= no eating alone |
| **Freezing** | **Vocalization** |
| 0= none | 0= frequent |
| 1= some episodes | 1= sometimes |
| 2= frequent episodes | 2= none |
